# A genome-wide association study for shared risk across major psychiatric disorders in a nation-wide birth cohort implicates fetal neurodevelopment as a key mediator

**DOI:** 10.1101/240911

**Authors:** Andrew J. Schork, Hyejung Won, Vivek Appadurai, Ron Nudel, Mike Gandal, Olivier Delaneau, David M. Hougaard, Marie Bækved-Hansen, Jonas Bybjerg-Grauholm, Marianne Giørtz Pedersen, Esben Agerbo, Carsten Bøcker Pedersen, Benjamin M. Neale, Mark J. Daly, Merete Nordentoft, Ole Mors, Anders D. Børglum, Preben Bo Mortensen, Alfonso Buil, Wesley K. Thompson, Daniel Geschwind, Thomas Werge

## Abstract

There is mounting evidence that seemingly diverse psychiatric disorders share genetic etiology, but the biological substrates mediating this overlap are not well characterized. Here, we leverage the unique iPSYCH study, a nationally representative cohort ascertained through clinical psychiatric diagnoses indicated in Danish national health registers. We confirm previous reports of individual and cross-disorder SNP-heritability for major psychiatric disorders and perform a cross-disorder genome-wide association study. We identify four novel genome-wide significant loci encompassing variants predicted to regulate genes expressed in radial glia and interneurons in the developing neocortex during midgestation. This epoch is supported by partitioning cross-disorder SNP-heritability which is enriched at regulatory chromatin active during fetal neurodevelopment. These findings indicate that dysregulation of genes that direct neurodevelopment by common genetic variants results in general liability for many later psychiatric outcomes.

## Introduction

Since the first attempts to establish a consistent nosology in psychiatry^1^ there has been an appreciation for the considerable difficulties in clearly delineating patient populations^2–4^. Without objectively discriminative biomarkers, diagnoses are made through the integration of subjective presentations, including patient experience and behavioral observations^2–4^. When research diagnostic systems are employed in clinically ascertained case-control studies, diagnoses may be designated hierarchically, incorporate perceived severity, censor milder episodes and/or select for archetypical presentations; a process that may obscure complicating comorbid or premorbid episodes^4–6^. In reality, psychiatric patients may present more heterogeneously, sharing features and blurring boundaries among disorders and between disorders and normal behavior^2,4,7^. To some extent this blurring is thought to extend to etiological features, with certain environmental risks predisposing to diverse outcomes^2,8,9^, nonspecific efficacy of drug treatments implying shared neurochemical pathologies^10^, and reports of extensive overlap in genetic risk factors^3,4^.

The large genetic component to susceptibility for psychiatric outcomes is established from consistently moderate to high heritability estimates^11^ and this contribution appears shared broadly among disorders. Initial observations of familial co-aggregation and genetic correlations for bipolar disorder and schizophrenia^12^ have been extended to include other mood disorders, autism spectrum disorder (ASD), and attention-deficit/hyperactivity disorder (ADHD)^5,13,14^. Recent SNP-based investigations of shared polygenetic risk provide further support in patient populations with similar symptom profiles, such as schizophrenia and bipolar^15–17^ or major depressive disorders^15,16,18^, but are also extending this overlap to disorders with more dissimilar clinical profiles such as schizophrenia and ADHD^16,19,20^, anorexia^16^ or ASD^15,21^. An emerging hypothesis is that at least a portion of genetic risk for psychiatric disorders is shared, or perhaps non-specific with respect to outcomes.

Plausible substrates for such non-specific susceptibility are emerging from molecular genetic studies. The pleiotropic effects of large effect copy number variants^5,22^ (CNVs) such as 16p11.2^23^, 22q11.2^24^ and NRXN1^25–27^ suggest insults to neurodevelopment and synaptic function may underlie some on the shared risk. The few studies of common variants directly investigating shared etiology^28–36^ have implicated genes involved in calcium channel neurobiology. A recent integrative transcriptional study implicated expression profiles of gene sets related to neuronal and astrocytic functions as shared molecular intermediaries^37^. Although plausible hypotheses are emerging, little has been done to characterize common variants that may have non-specific effects, especially in more representative patient populations.

Here we leverage the unique Integrative Psychiatric Research Consortium (iPSYCH) case-cohort study^38^. iPSYCH is composed of one of the largest single population samples of genotyped psychiatric patients in the world and a diagnosis-free, random sample from the same birth cohort. iPSYCH has the unique advantage of being constructed from a single population and diagnostic schema, uniformly ascertained according to care provided under the same public healthcare system. By comparison, the majority of psychiatric disorder cohorts are ascertained using different schemas and in different populations which may confound, obscure, or otherwise diminish cross-disorder inferences.

We performed a genetic dissection of this naturalistic and essentially complete patient population ascertained passively from a nation-wide Danish birth cohort (1981–2005, N=1,472,762) for attention-deficit/hyperactivity disorder (ADHD), affective disorder (AFF), anorexia (ANO), autism spectrum disorder (ASD), bipolar disorder (BIP) or schizophrenia (SCZ; Supplementary Table 1,2; Supplementary Figure 1). We perform a diagnosis agnostic, crossdisorder (XDX) GWAS comparing common variant allele frequencies between the entire patient (N=46,008) and population control cohorts (N=19,526). We explore the consistency of our findings in published reports and perform an independent replication study in a small, diverse sample from the same birth cohort (N=7,163). Finally, we integrate published neurobiological data to suppose that the effects of common genetic variants with non-specific associations to psychiatric outcomes are mediated through dysregulation of early neurodevelopmental processes.

**Table 1.**
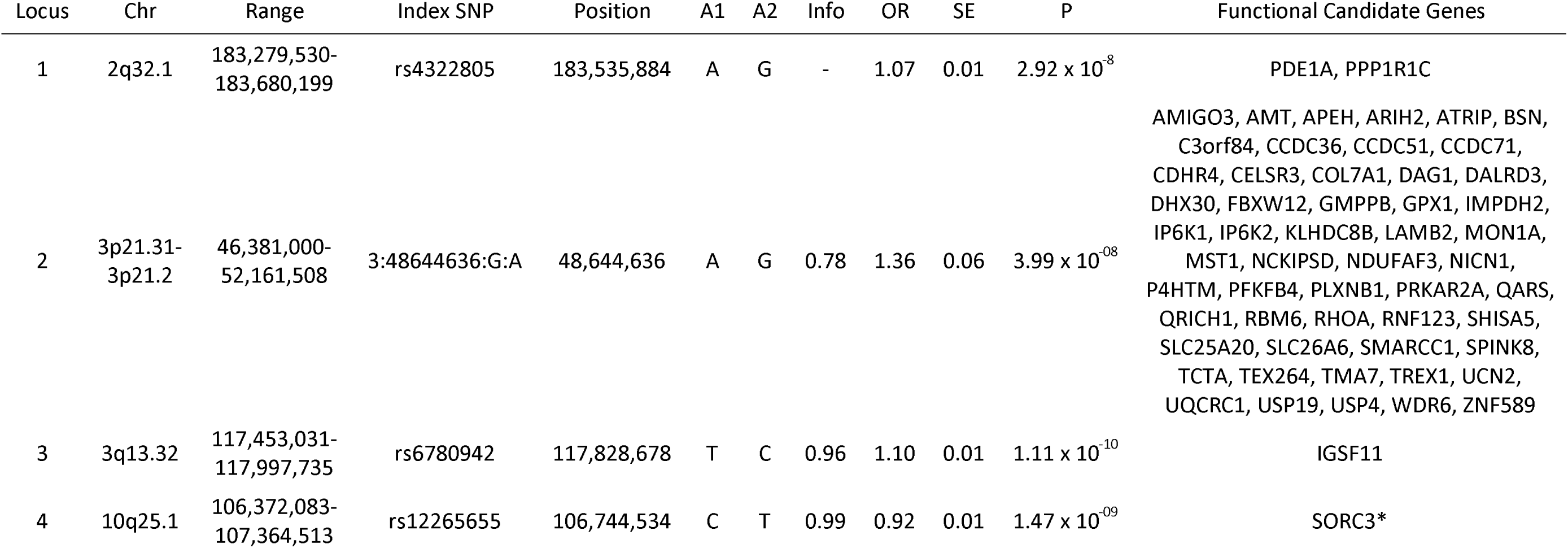
Genome-wide Significant Associations. Four loci indexed by genome-wide significant index SNPs in the XDX GWAS implicate a number of candidate genes. Standard Errors are for logistic regression coefficients and this on the natural log scale (In(OR)). OR, Odds Ratio; SE, Standard Error; Al, Effect Allele; A2, Non-Effect Allele; Info, Imputation Information Score; Z, Regression coefficient test statistic; P, p-value. *SORCS3 was implicated by overlap, not functional connection.

**Table 2.** Independent Replication Support. Where data was available, the trends for each locus are generally consistent in a number of external sources. *Significant after correction for multiple testing. ^**^Additionally, locus 4 has recently been reported as genome-wide significant (p < 5×10^−8^) in two studies including iPSYCH^18,20^. OCD-ASD, Joint Obsessive-Compulsive Disorder -Autism Spectrum Disorder.

| Locus | Disc | Repl |  | NHGRI GWAS Catalog |  |  | eADHD |  | eAFF |  | eANO |  | eASD |  | eBIP |  | eSR-MDD |  | eSCZ |  | eXDX |  |
| --- | --- | --- | --- | --- | --- | --- | --- | --- | --- | --- | --- | --- | --- | --- | --- | --- | --- | --- | --- | --- | --- | --- |
|  | Sign | Sign | P | Sign | P | Trait | Sign | P | Sign | P | Sign | P | Sign | P | Sign | P | Sign | P | Sign | P | Sign | P |
| 1 | + | + | 0.26 | NA | NA | NA | - | 0.81 | - | 0.63 | - | 0.96 | - | 0.56 | - | 0.58 | NA | NA | + | 0.14 | - | 0.52 |
| 2 | + | + | 0.14 | + | $7 \times 10^{-6}$ | OCD-ASD | + | $1 \times 10^{-3}$ | + | 0.05 | + | 0.09 | + | 0.07 | + | 0.01 | NA | NA | + | $9 \times 10^{-4}$ | + | $3 \times 10^{-3}$ |
| 3 | + | + | 0.3 | NA | NA | NA | NA | NA | NA | NA | + | 0.35 | + | 0.28 | + | 0.26 | NA | NA | + | 0.18 | NA | NA |
| 4** | - | - | 0.16 | - | $2 \times 10^{-6}$ | SCZ | + | 0.92 | - | 0.05 | - | 0.06 | - | 0.3 | - | 0.43 | - | $1 \times 10^{-5}$ | - | $8 \times 10^{-5}$ | - | 0.02 |

**Figure 1.**
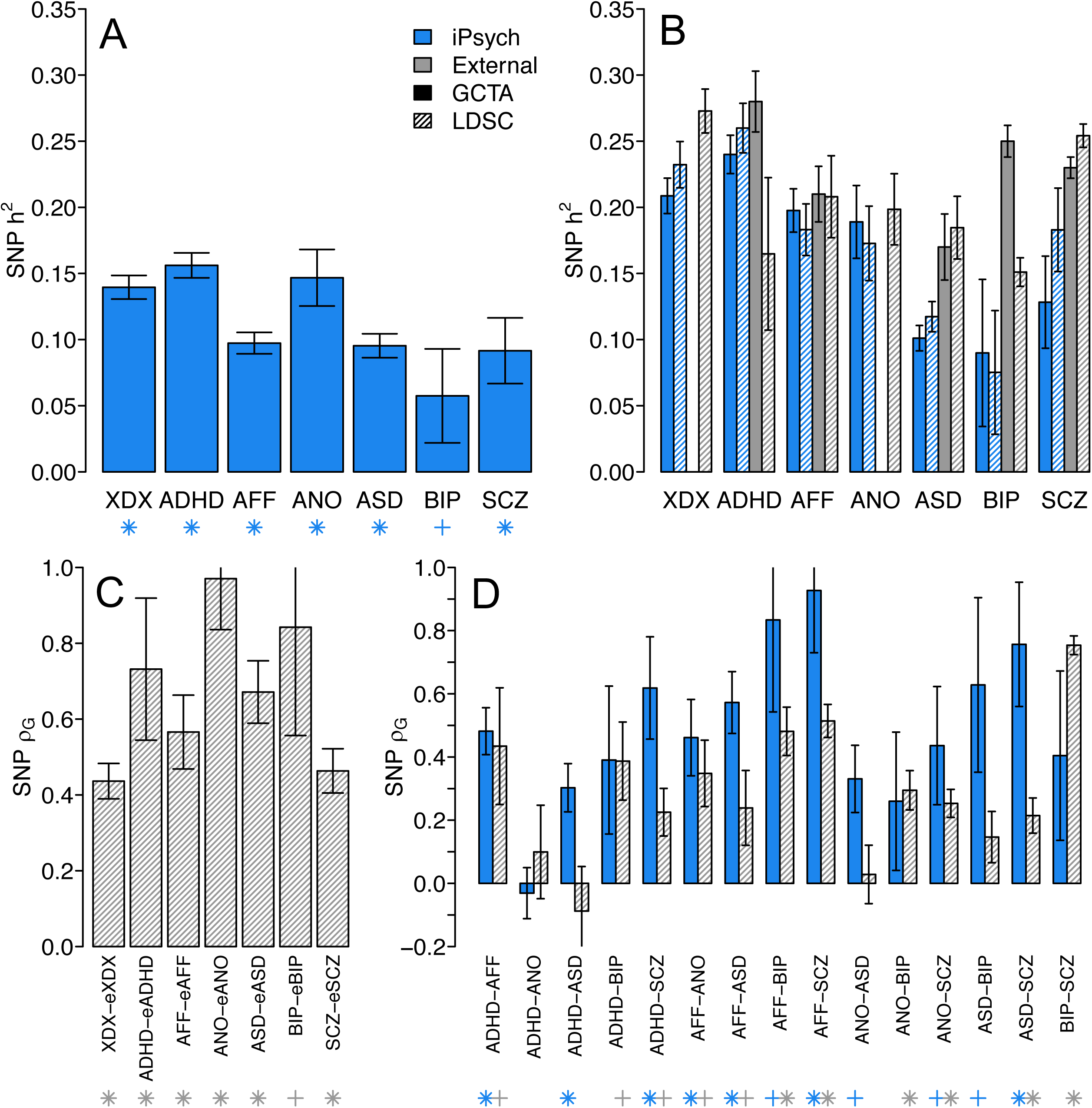
SNP heritability and genetic correlation estimates for iPSYCH indications. (A) SNP-heritability estimates are transformed to the liability scale defined by prevalence in the random population sample (XDX, 0.088; ADHD, 0.010; AFF, 0.013; ANO, 0.003; ASD, 0.008; BIP, 0.001; SCZ, 0.002). (B) SNP-heritability estimates for each iPSYCH indication (blue bars) are estimated via GREML (solid bars) and LDSC (striped bars) and presented in comparison with estimates from representative external studies (solid and striped grey bars). Estimates here are presented on the same liability scale defined by typical prevalence estimates (XDX, 0.35; ADHD, 0.05; AFF, 0.15; ANO, 0.01; ASD, 0.01; BIP, 0.01; SCZ, 0.01) (C) The SNP-based genetic correlations between iPSYCH and external studies were estimated via LDSC. (D) SNP-based genetic correlations between each iPSYCH indication estimated via GREML (solid blue bars) are presented beside correlations estimated from external studies via LDSC (grey bars). XDX, crossdiagnosis; ADHD, attention-deficit, hyperactivity disorder; AFF, affective disorder; ANO, anorexia; ASD, autism-spectrum disorder; BIP, bipolar disorder; SCZ, schizophrenia; OTH, other. Star, p < 5 × 10^−4^; Cross, p < 0.05. Error bars denote standard errors.

## Results

### The Genetic Architecture of iPSYCH Diagnostic Indications

The SNP-heritability for and genetic correlations among indications in the iPSYCH patient population are depicted in Figure 1 (Supplementary Tables 3–8). All estimates were highly significant (p < 5 × 10^−4^) except for the small BIP cohort (p = 0.049). The SNP-heritability for diagnosis agnostic psychiatric indications (XDX) was of similar magnitude to that for each individual disorder (Figure 1A, Supplementary Table 3). We compare these estimates to those taken from external studies for the same disorders (Figure 1B, Supplementary Table 4), including linkage disequilibrium score regression (LDSC) estimates in seven external GWAS: attention-deficit hyperactivity disorder^39^ (eADHD), autism spectrum disorder^40^ (eASD), major depressive disorder^41^ (eAFF), eating disorders^42^ (eANO), bipolar disorder^43^ (eBIP), schizophrenia^44^ (eSCZ) and a cross-disorder (eXDX) GWAS^34^ combining portions of eADHD, eAFF, eASD, eBIP and eSCZ. SNP-heritability estimates by GREML and LDSC for the iPSYCH indications are broadly similar to each other and to those from external studies when presented on the same liability scale, although differences are present for adult onset disorders (SCZ and BIP) and ASD. LDSC standard errors are consistent with highly significant estimates, with the exception of iPSYCH BIP and eADHD. LDSC estimates of genetic correlation between iPSYCH indications and external GWAS of the same disorders were also highly significant (p < 5 × 10^−4^), with only BIP showing nominal significance (p = 3.2×10^−3^; Figure 1B, Supplementary Table 5).

GREML point estimates of genetic correlations for all pairs of iPSYCH indications are at least moderate (ρ_G_ > 0.25; Figure 1C, Supplementary Table 6–7), with the exception of ADHD-ANO. Among the iPSYCH indication pairs, seven produced highly significant estimates (p < 5 × 10^−4^; ADHD-AFF, ADHD-ASD, ADHD-SCZ, AFF-ANO, AFF-ASD, AFF-SCZ, and ASD-SCZ) and four were nominally significant (5 × 10^−4^ < p < 0.05; AFF-BIP, ANO-ASD, ANO-SCZ, ASD-BIP). Only ADHD-ANO, and three involving the smallest case group, BIP, were not significant (p > 0.05; ADHD-BIP, ANO-BIP, and BIP-SCZ). LDSC genetic correlation estimates among external GWAS (Figure 1C; Supplementary Table 8) are nearly consistently smaller than for iPSYCH indications of the same disorders, with eBIP-eSCZ and non-significant eADHD-eANO the only exceptions. Among the external LDSC estimates eAFF-eBIP, eAFF-eSCZ, eANO-eBIP, eANO-eSCZ, eASD-eSCZ, eBIP-eSCZ were highly significant (p < 5×10^−4^), eADHD-eBIP, eADHD-eAFF, eADHD-eSCZ, eAFF-eANO, eAFF-eASD were nominally significant (5×10^~4^ < p < 0.05) and eADHD-eANO, eADHD-eASD, eANO-eASD and eASD-eBIP were not significant (p > 0.05). The genetic architecture of psychiatric disorders implied by iPSYCH register indications appears qualitatively similar to that presented in the previous reports based on different ascertainment schemes^16^'^45^, although the iPSYCH genetic correlations are generally larger.

### A Diagnosis Agnostic GWAS

Motivated by the appreciable XDX SNP-heritability and prevalence of substantial genetic correlations among iPSYCH indications, we performed a diagnosis agnostic, cross-disorder GWAS (XDX). We combined all iPSYCH psychiatric patients into a single case cohort (N=46,008) and used a diagnosis-free, random population sample as controls (N=19,526). We identified four independent loci tagged by genome-wide significant index SNPs (referred to throughout as loci 1–4; p < 5×10^−8^; Table 1; Supplementary Figure 2) and another 46 loci (loci 5–50) indexed by suggestive associations (stratified false discovery rate, sFDR, < 0.05; Supplementary Table 9). No locus showed genome-wide significant evidence of a second association. The distribution of p-values from the GWAS demonstrated moderate levels of inflation (genomic inflation factor^46^, A,gc=1–15; Supplemental Figure 3). A significant LDSC estimate for SNP-heritability (observed scale h^2^=0.13; s.e.=0.01) with an intercept close to unity (1.02; s.e.=0.01) suggests this inflation is likely due to polygenes rather than population stratification or cryptic relatedness.

**Figure 2.**
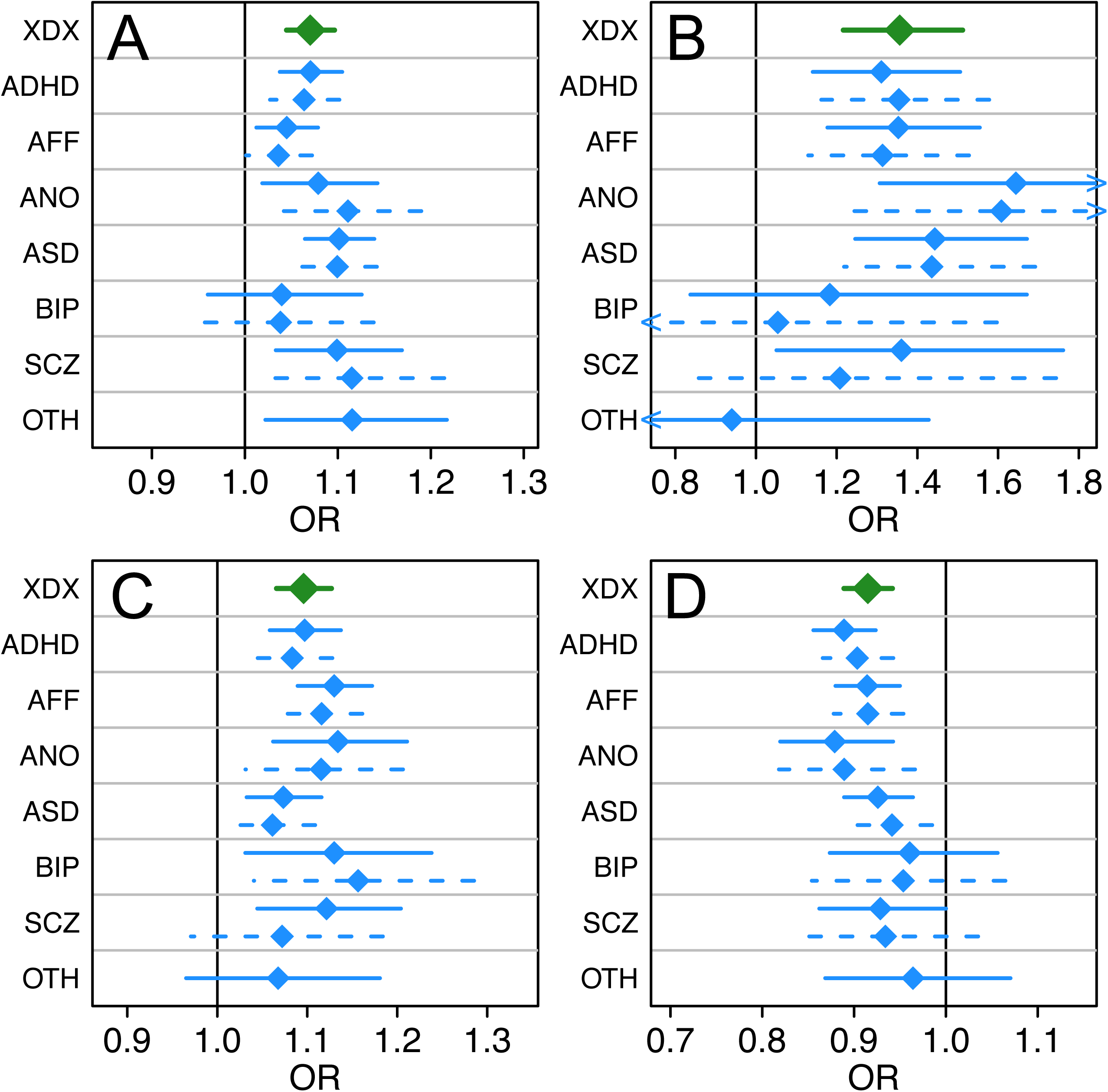
Odds ratio estimates for single indications. For each of locus 1 (A), locus 2 (B), locus 3 (C), and locus 4 (D), the odds ratio (OR) and approximate 95% confidence interval (CI) from the XDX GWAS (green diamond and solid bar) is presented with the estimates for each single indication. Odds ratios were estimated using all cases for each indication (blue diamond and solid bars) and restricting to cases without co-occurring indications (blue diamond and dashed bars). Odds ratios are also estimated for the patients with no ascertained indications, but at least one other psychiatric indication (OTH). Although generally consistent, the smaller groups (SCZ, BIP, OTH) are, as expected, less informative (largest CIs). XDX, cross-diagnosis; ADHD, attention-deficit, hyperactivity disorder; AFF, affective disorder; ANO, anorexia; ASD, autism-spectrum disorder; BIP, bipolar disorder; SCZ, schizophrenia; OTH, other.

**Figure 3.**
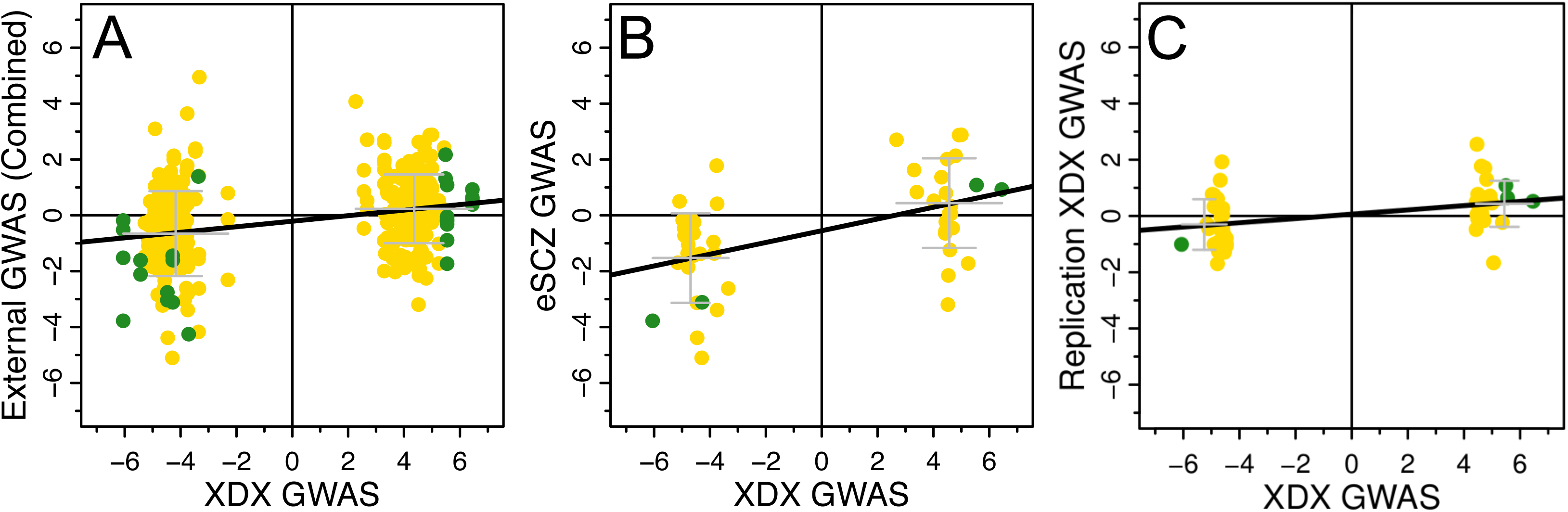
Consistency of Effects in Independent Studies. The z-scores for the best proxy SNPs in independent GWAS are plotted against the z-scores for the same SNPs from the XDX GWAS. Figures shows XDX effects predicting (A) eADHD, eAFF, eANO, eASD, eBIP, eSCZ, eSR-MDD, and eXDX in aggregation, (B) eSCZ only, and (C) iPSYCH replication cohort effects. Genome-wide significant XDX loci 1–4 are shown with green dots and suggestive loci 5–46 are shown with gold. Black lines show the regression fit. Grey bars depict the mean and one standard deviation interval of replication effects for SNPs with positive or negative effects in the XDX GWAS. All regression fits are significant (p < 5 × 10^−3^; All: p = 6.5×10^−9^; eSCZ: p = 1.3×10^−4^; iPSYCH Replication: p = 4.2×10^−3^).

Although the XDX GWAS is most sensitive to variants with consistent effects across diagnoses, variants strongly associated with a single indication could present as apparent crossdisorder associations. However, rather than showing skewing of the odds ratios for one or two indications, the index SNPs for loci 1–4 show consistent trends across nearly all indications when association tests were performed using one-indication case groups (Figure 2). This broad pattern holds for suggestive loci 5–50 (Supplementary Tables 10^−^11). The XDX GWAS uncovered multiple loci with non-specific effects on risk across iPSYCH indications that are not driven by a single or small subset of disorders.

### Independent Statistical Support

We sought independent statistical support for our associations from three sources. First, we queried the top associations from 3,893 GWAS aggregated in the 13 November 2017 release of the NHGRI-EBI GWAS catalog^47^ for connections to our top 50 XDX loci (Table 2; Supplementary Table 17). The catalog ontology places psychiatric outcomes within the broader label of “neurological disease.” In total, our 50 loci were related to 18 out of 539 neurological disease GWAS and 17 out of 3,354 other traits and diseases (Table 2, Supplementary Table 12), a significant difference (binomial proportion test; p = 5×10^−11^). Among XDX loci 1–4, locus 2 showed significant independent support (p < 2.32×10^~5^ correcting for four loci, each potentially connected to 539 GWAS) in a joint obsessive-compulsive disorder (OCD)-ASD GWAS^48^ and locus 4 showed support in a schizophrenia GWAS^49^. Locus 4 was recently described as genome-wide significant in two meta-GWAS that include the iPsych data but were not in the GWAS catalog at the time of writing, one for MDD^18^ and one for ADHD^20^. Among the 46 suggestive loci, locus 9 and locus 36 had significant independent support (p < 1.86×10 ^6^ correcting for 50 loci and 539 GWAS) among genome-wide significant loci for schizophrenia^44^. Locus 23 was supported by an association with depressive symptoms^50^. A few suggestive loci had significant evidence for potential pleiotropy (p < 2.98×10^~7^ correcting for 50 loci and 3,354 GWAS; Supplementary Table 12). Locus 9 was additionally associated with aneurysm^51,52^ and LDL cholesterol levels^53^, locus 13 was associated with age at menarche^54^ and a hematologic measurement^55^, locus 33 was provocatively associated with vitamin B12^56,57^ and homocysteine^58^ levels, and locus 50 was associated with vitiligo^59^.

The GWAS catalog only contains a sparse selection of SNPs from the studies it summarizes which limits the opportunity for comprehensive replication and aggregate concordance testing. Thus, as a second external validation, we mapped our top 50 loci to the best proxy SNP in each of the seven external GWAS described previously (eADHD, eAFF, eANO, eASD, eBIP, eSCZ, eXDX) and, where possible, to the best proxy among the top 10,000 associations from a self-report major depressive disorder GWAS^60^ (eSR-MDD; Table 2, Supplementary Table 13). Doing so, we observe significant external replication (p < 1.56×10^−3^ correcting for four loci, each tested in eight studies) for locus 2 in the eADHD and eSCZ studies. Locus 4 significantly replicated in the eSR-MDD and eSCZ studies. Among the suggestive loci 5–50, we observe significant replication (p < 1.25×10^−4^ correcting for 50 loci and eight studies) for locus 9 and locus 11 in the eSR-MDD GWAS. Even where not individually significant, the association trends are strikingly consistent across all studies for loci 2–4. Importantly, we the XDX trend is consistent for each of loci 1–4, and 35 out of 46 suggestive loci, in the eSCZ GWAS. As a test of this concordance, we used linear regression to ask if, across the top 50 loci, the signed effects (regression statistics; z-scores) for proxy SNPs in each external GWAS, as well their combination, are significantly predicted from the XDX effects for the same SNPs. We observed significant concordance (p < 5 × 10^−3^ to correct for ten concordance tests) between the XDX effects and the aggregated effects from all seven external GWAS (Figure 3A) as well as in the eSCZ GWAS alone (Figure 3B). Concordance was also significant for the eADHD and eXDX GWAS (Supplementary Figures 4–10).

**Figure 4.**
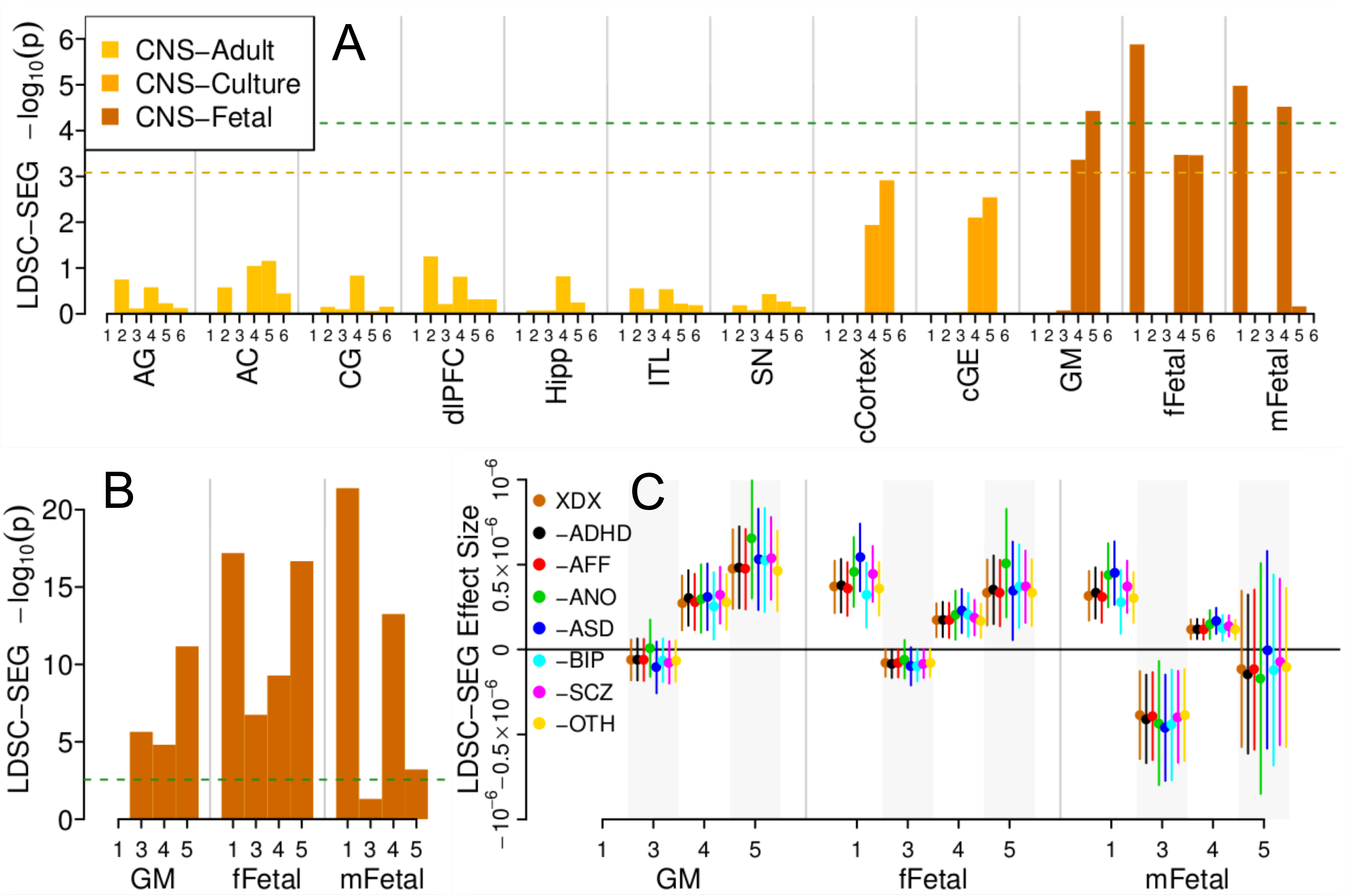
LDSC Partitioning. (A) Each bar depicts the -logio(p-value) from an LDSC-SEG test of enriched XDX GWAS heritability at variants annotated with different chromatin marks in different tissues. Dashed lines represent significant (green; p < 8.32×10^−5^) and suggestive (gold; FDR < 0.05) enrichment. (B) Enrichment p-values using the same fetal brain annotations in the eSCZ GWAS. Green dashed bars denote significant enrichment (p < 2.78×10^−3^). (C) Original enrichment effect sizes (orange circle and 95% confidence interval) are shown to be similar to re-estimates computed by iteratively excluding all patients with a given indication. CNS, central nervous system; 1, DNAse; 2, H3K27ac; 3, H3K36me3; 4, H3K4me1; 5, H3K4me3; 6, H3K9ac. XDX, cross-disorder; eSCZ, external schizophrenia; AG, Adult Angular Gyrus; AC, Adult Anterior Caudate; CG, Adult Cingulate Gyrus; dIPFC, Adult Dorsolateral Prefrontal Cortex; Hipp, Adult Hippocampus; ITL, Adult Inferior Temporal Lobe; SN, Adult Substantia Nigra; cCortex, Cortex Derived Cultured Neurospheres; cGE, Ganglion Eminence Derived Cultured Neurospheres; GM, Gemrinal Matrix; fFetal, female Fetal Brain; mFetal, male Fetal Brain.

As a third form of external validation, we repeated the association tests for each index SNP using linear mixed models in a small, independent replication cohort. We included subjects excluded from the discovery GWAS cohort on the basis of outlying genetic ancestry (replication cohort: N=7,163, 4,481 cases; Supplementary Table 1; results: Table 2, Supplementary Table 14). Given the reduced sample size we did not expect, nor did we observe, any significant individual replications. We did observe, perhaps surprisingly given the diverse genetic backgrounds, a consistent trend at each of loci 1–4 and for 31 of the additional 46 suggestive loci. This broad concordance was also significant by the same linear model as above (p < 5.56×10^−3^; Figure 3C). Taken together, these analyses provide several forms of independent statistical support for the XDX loci identified here.

### Tissue Specific Partitioning of Non-Specific Psychiatric Diagnosis Heritability

To determine if XDX loci capture a coherent biological process, we first partitioned the XDX SNP-heritability among tissues. We used LDSC regression for specifically expressed genes^61^ (LD-SEG) to test for significant enrichments in average per-SNP heritability across 601 variant annotations (LD scores) related to genes preferentially expressed or chromatin open and active in various tissues. None of the expression annotations (GTEx and DEPICT LD scores) showed even suggestive evidence (FDR > 0.05; Supplementary Figures 11–12, Supplementary Table 15-16) for enrichment. Among the RoadMap chromatin annotations (Figure 4A, Supplementary Table 17), several features related to gene regulation in fetal brain showed significant (p < 6.88×10^−5^; germinal matrix: H3K4me3, p=3.73×10^−5^; female fetal brain: DNase, p=1.31×10^−6^; male fetal brain: DNase, p=1.05×10^−5^, H3K4me1, p=3.02×10^−5^) or suggestive (FDR < 0.05; germinal matrix: H3K4me1, p=4.34×10^−4^; female fetal brain: H3K4me1, p=3.38×10^−4^, H3K4me3, p=3.45×10^−4^) enrichment. No significant or suggestive enrichment was observed for non-brain RoadMap LD-scores (Supplementary Figure 13, Supplementary Table 18). Enrichment in the same fetal brain chromatin annotations replicated in the eSCZ GWAS, a result previously described in the LDSC-SEG report^61^ (Figure 4B, Supplementary Table 19). Furthermore, iteratively removing subjects with each indication from the iPSYCH cases and re-estimating enrichment produced consistent results (Figure 4C, Supplementary Table 20). Figures 2B and 2C emphasize the disorder non-specificity of these variant enrichments. The heritability enrichments for regulatory variants active in fetal brain seen for the XDX GWAS replicate as a class in an independent psychiatric GWAS for a disorder (SCZ) among the least represented in the iPSYCH patient population (Supplementary Figure 1) and are not driven by any single iPSYCH indication.

### Identifying and Characterizing Candidate Genes

We used FineMap^62^ at each locus to identify a set of credible SNPs, which were subsequently connected to candidate target genes using chromatin interaction maps for the developing cortical laminae, as previously described^63^. Out of 627 credible SNPs, 400 implicated one or more of 281 candidate genes (160 protein coding; Supplementary Table 21). As a set, these genes were enriched for genes that regulate axon and dendrite development, receptor Tyrosine kinases, and histone regulators (Figure 5A). Notably, histone regulators and chromatin remodelers have been recently implicated in various psychiatric disorders, including ASD and SCZ^21,64,65^. Consistent with the heritability enrichment, the candidate genes were more highly expressed during prenatal stages (p = 1.67×10^−10^, Figure 5B), with peak expression during mid-gestation in the human brain (Figure 5C). To understand what cells these genes might be functioning in, we leveraged expression profiles of single cells extracted from fetal brain tissue and tested for enrichment.^66^ Remarkably, we observe candidates to be enriched for genes preferentially expressed in radial glia, an early form of neural stem cell, and interneurons (FDR < 0.05; Figure 5D). These results suggest that the cross-diagnosis loci are implicated in the proliferation of neural stem cells and establishment of neural circuits during brain development.

**Figure 5.**
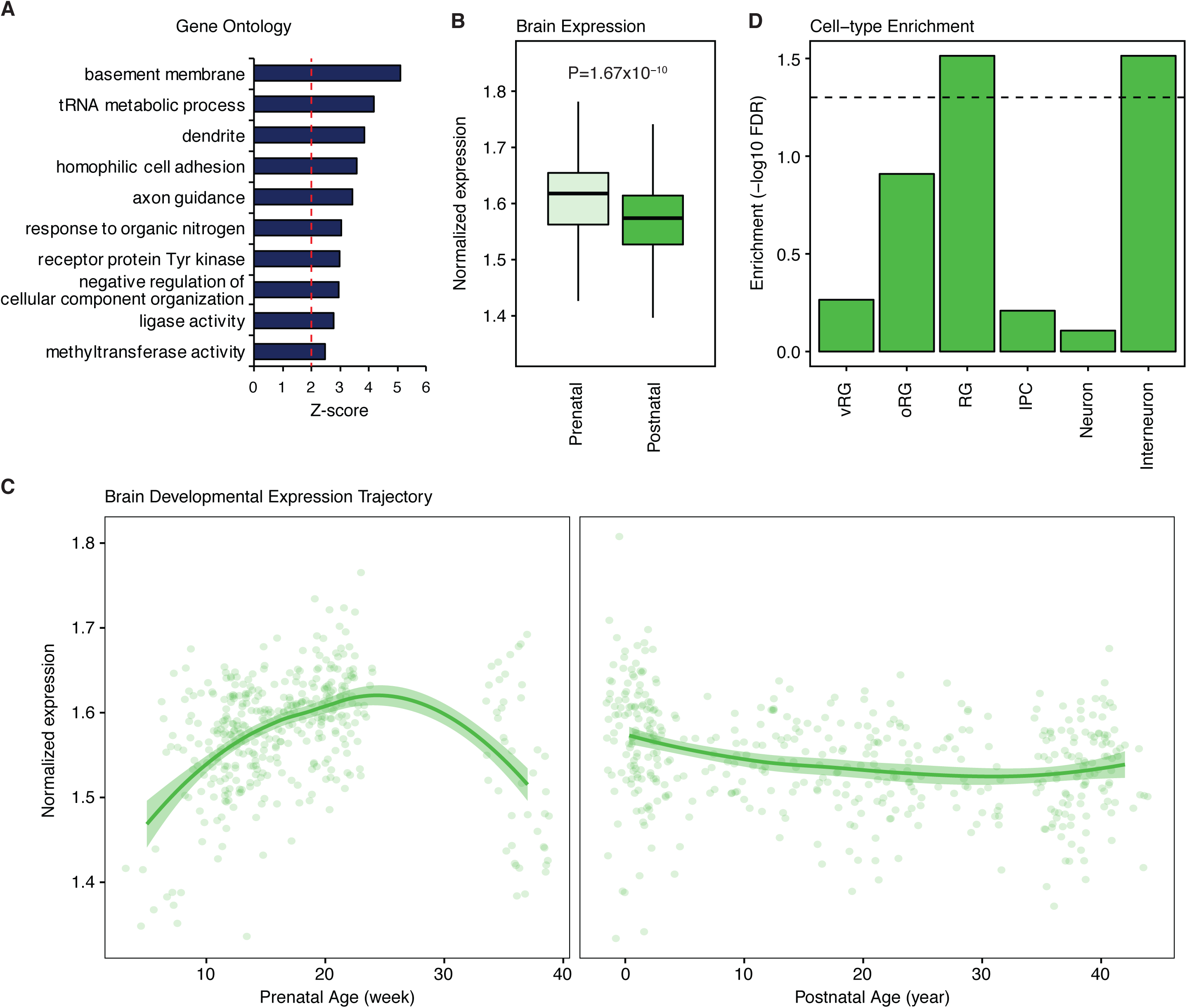
Candidate Gene Set Enrichments for Neurodevelopmental Processes. (A) Top ten gene ontology categories significantly enriched among the 281 XDX candidate genes. (B) The XDX candidate genes have higher average cortical expression in the prenatal stage. (C) The average cortical expression trajectory for XDX candidate genes peaks during midgestation. (D) The XDX candidate genes are enriched for genes with specific expression in developing radial glia and interneurons. XDX, cross-diagnosis. RG, radial glia; vRG, ventricular radial glia; oRG, outer radial glia; IPC, intermediate progenitor cells.

Specific candidate genes implicated by our significant loci include PDE1A and PPP1R1C for locus 1 (annotated local Manhattan plot: Supplementary Figure 14). PDE1A is a Ca2+/calmodulin dependent cyclic nucleotide phosphodiesterase (PDE1s) that regulates cAMP and cGMP^67^ and can regulate L-type and T-type voltage gated calcium channels^68^. PPP1R1C has been shown to regulate neurite growth in cultured neurons^69^ and is an inhibitor of protein phosphatase 1 (PP1), a protein involved in synaptic transmission and plasticity as part of the post synaptic density^70^. Locus 2 is indexed by an uncommon SNP (minor allele frequency = 0.018) and covers a broad, gene dense region implicating 53 diverse genes (Supplementary Figure 15–16). Among these, DAG1, QRICH1, RNF123 and SMARCC1 harbor *de novo* risk variants for ASD and CELSR3 for SCZ^71,72^. Locus 3 implicates IGSF11 (Supplementary Figure 17), a neuronal adhesion molecule that binds to and stabilizes AMPA receptors regulating synapse development and plasticity^73^. Locus 4 overlaps considerably with the body of SORC3 (Supplementary Figure 18), which encodes a postsynaptic sorting and signaling receptor involved in aspects of neuronal functioning including synaptic plasticity^74,75^.

Among the suggestive loci, candidate genes implicated by locus 14 (WHSC1), locus 27 (TENM2), locus 31 (THSD7A) and locus 47 (CDH13) also harbor *de novo* variants for ASD and locus 36 (ZDHHC5) for SCZ^71–72^. Locus 16 implicates LPHN3, a gene that regulates synaptic connectivity between cortical layers 2/3 and 5^76^, and locus 29 implicates ELFN1, a gene involved in the establishment of somatostatin-positive interneuronal circuits^77^; two exemplars of the developmental interneuronal enrichment. Interestingly, both locus 32 (Supplementary Figure 19) and locus 33 (Supplementary Figure 20) implicate CUBN, a multi-ligand endocytic receptor involved in the absorption of vitamin B12^78^, and Imerslund-Gräsbeck Syndrome^79^, an inherited disorder of vitamin B12 malabsorption. Brief refSeq^80^ and UniProt^81^ descriptions for all candidate protein coding genes were aggregated from UCSC genome browser databases^82,83^ and collated in Supplementary Table 22.

## Discussion

In this study, we leveraged the uniquely designed iPSYCH study to provide an unprecedented perspective on the overlap in genetic etiology underlying major psychiatric disorders. The iPSYCH cohort has the advantage of coming from a nationally representative study interrogating an essentially complete population of patients against a sample of diagnosis free individuals from the same Danish birth cohort. Previous results are based on cohorts identified independently at tertiary research centers by different research groups, often in different countries, factors which could reduce true XDX overlap, or shared heritability. Here, patients are ascertained against a uniform background, including genetic background, diagnostic schema, health care system and other cultural or sociodemographic factors, limiting the potential impact of heterogeneity that may arise when comparing independently ascertained patient cohorts.

The picture of the genetic architecture inferred from the iPSYCH indications is similar to that painted by previous studies, with a few possible exceptions. First, we observe slightly less SNP-heritability for adult onset disorders (BIP and SCZ) in iPSYCH. We note that these are among the smallest patient cohorts and for disorders that, given the relative youth of the targeted birth cohort (1981–2005, mean age 20.4), are likely not yet fully expressed. Although this feature could lead us to underestimate disorder specific heritability, it is a strength when interpreting the external consistency of the heritability partitioning and single variant associations because a large schizophrenia GWAS would be less likely to replicate iPSYCH disorder specific variants, those that would be implicated by the largest iPSYCH cohorts of ASD, ADHD and AFF. The replication of our results in the eSCZ study, in particular, then makes their plausibility as non-specific findings that much greater.

We also observe consistently larger genetic correlations between disorders in iPSYCH. There are multiple potential explanations for this including the reduced background heterogeneity, comorbidity, or diagnostic misspecification^84^. The ascertainment schema iPSYCH employs, passively aggregating health register indications absent an integrative clinical research assessment at the time of collection, could then be perceived as a weakness. However, it is neither obvious how to best approximate an integrative research diagnosis from aggregated register indications algorithmically, nor that that represents an optimal goal. The representativeness of the iPSYCH patients uniquely exemplify a target for precision medicine based initiatives. Approximating a more archetypical research ascertainment would lead us to alter complicating comorbid or premorbid diagnostic indications or censor difficult to classify patients. One could argue this would potentially move us further from the populations we aim to understand. Rather than deviate from the strength of this cohort, we present the results as a bridge towards a more representative assessment of genetic overlap in a naturalistic population of patients with major psychiatric diagnoses. We also see this as motivation for future investigations into the role that diagnostic schema play in shaping our conceptualizations of these disorders. With the continued emergence of large population-based ascertainments, such as iPSYCH, 23andMe^60^, the UKBiobank^85^, and large insurance record cohorts^86^'^87^, we feel this point is critically important for understanding potential differences so we may integrate inferences from diverse study designs rather than censor them away.

It is interesting that our functional interpretations of shared genetic risk factors, both in general and with respect to our top associations, point to fetal neurodevelopmental processes. This epoch has been implicated by previous studies of large effect CNVs and rare or *de novo* variants, but also of environmental exposures^88,89^. Such convergence could suggest a critical developmental window during which part of the susceptibility to later psychiatric outcomes is defined. The overlap in timing for the action of genetic and environmental susceptibility factors may also help to carve out a hypothesis space for targeted investigations of gene-environment interactions.

While germline variants are present since the first moments of embryonic development, the chromatin they reside in and genes they may target undergo a continual evolution of active and inactive states across developmental and life stages. We note that our top associations are potential regulators of genes with familiar functions^45–65^, (post)synaptic and calcium channel biology. We also demonstrated that, in our study, these variants and their associated genes are most coherently involved in aspects of fetal neurodevelopment. If susceptibility emerges from disruptions to this familiar synapse and calcium channel biology but specifically during neuronal proliferation, migration and establishment of circuits, this could have implications for the development of interventions. The pathology induced by these genes may not be directly treatable by pharmacological treatments administered during much later developmental stages when symptoms present. To make good on the promise that GWAS can identify plausible pharmacological targets, it will become critical to consider the developmental stages during which variants induce susceptibility. It may be the case that the pathology induced by an associated germline variant occurs in one developmental stage, but actionable pharmacological targets represent *different* downstream molecular pathologies^37^. Explicitly considering the developmental course of pathological susceptibility implied by associated germline variants is a critical next step in the translational promise of GWAS results and emphasizes the importance of partitioning risk with development in mind.

One unexpected suggestive finding to emerge from this work was potential pleiotropy between psychiatric outcomes and vitamin B12 metabolism through CUBN. There is a long literature implicating vitamin B12 and related pathways in psychiatric outcomes^90–93^, however, no studies we are aware of have implicated common genetic variants related to CUBN as a contributing factor. Future work should confirm this finding and more definitively characterize potential biological consequences of variants within this locus. Ideally one could test if it represents mediated pleiotropy, whereby these variants alter vitamin B12 metabolism leading to deficiency and subsequent deficiency related psychiatric outcomes, true pleiotropy through a parallel function of the gene, or false pleiotropy through undescribed linkage.

Even though our sample is large for a single psychiatric cohort, it is still comparatively modest when contrasted with what could be needed to fully characterize the specificity of loci for single disorders and subsets of disorders, especially under our analytic design. Thus, it is also likely that other susceptibility factors are more specific, contributing to the unique presentations of each disorder. Future work could take direct aim at these factors as we continue to update, partition and functionally characterize our conceptualizations of the genetic architecture underlying major psychiatric disorders.

## Methods

### The iPSYCH Study Cohort and Data

The Integrative Psychiatric Research Consortium (iPSYCH) cohort is described in detail elsewhere^38^. Briefly, iPSYCH samples from the entire population of Denmark born between 1981 and 2005 (N=1,472,762). In total, 87,764 individuals were selected. 30,000 individuals were sampled randomly without regard for psychiatric disorders. The remaining 57,764 were ascertained for indications of clinical diagnoses recorded in the Danish Civil^94^, National Patient^95^ and/or Psychiatric Central Research^96^ Registers describing care received for attention-deficit/hyperactivity disorder (ADHD), anorexia (ANO), autism spectrum disorder (ASD), affective disorder (AFF), bipolar affective disorder (BIP), or schizophrenia (SCZ). Where available a dried blood spot was obtained from the Danish Neonatal Screening Biobank^97^.

For this study, indications were from the Danish National Patient Register^95^ (complete through 2012) and Psychiatric Central Research Register^96^ (PCRR; complete through 2013), and coded according to criteria previously described^98^. Indications are International Classification of Disease^99,100^ (ICD) codes representing the clinical diagnosis associated with an instance of care provided at one of many psychiatric facilities throughout Denmark. Records include all admissions to inpatient psychiatric facilities since 1969 and outpatient psychiatric care received in all psychiatric hospitals, psychiatric wards and emergency rooms since 1995^96^. Care administered by primary care or private practice physicians is not recorded into national registers. Diagnoses prior to 1994 were associated with codes from the ICD 8^th^ revision^100^ and converted to equivalent ICD 10^th^ revision^99^ (ICD10) codes to match the majority of indications^98^. We define the following seven cases groups as having at least one indication with the corresponding ICD10 (or equivalent ICD8) codes: cross-diagnosis (XDX: F00–99), attention-deficit/hyperactivity disorder (ADHD: F90.0), affective disorders (AFF: F32-F39), anorexia (ANO: F50.0, F50.1), autism spectrum disorders (ASD: F84.0, F84.1, F84.5, F84.8, F84.9), bipolar disorder (BIP: F30–31), and schizophrenia (SCZ: F20). As an additional case group, we consider those patients with other psychiatric indications, exclusively (OTH: F00-F99 not listed previously, only). Case status can reflect the presence of a single or multiple indications and were not censored or integrated hierarchically. The counts, prevalence and co-occurrence of indications are shown in Supplementary Tables 1–2 and co-occurrence is depicted in Supplementary Figure 1.

Initial genotyping was performed at the Broad Institute with amplified DNA extracted from dried blood spots and assayed on the Inifinium PsychChip vl.O array^38^. In total, 77,639 subjects were successfully genotyped across 25 waves at ~550,000 SNPs. A subset of good quality common SNPs (N=246,369) were phased into haplotypes in a single batch using SHAPTEIT3^101^ and imputed in 10 batches using Impute2^102^ with reference haplotypes from the 1000 genomes project phase 3^103^. Imputed additive genotype dosages and best guess genotypes were checked for imputation quality (INFO > 0.2), deviations from hardy Weinberg equilibrium (HWE; p < 1×10^−6^), association with genotyping wave (p < 5 × 10^−8^), association with imputation batch (p < 5 × 10^−8^), and differing imputation quality between cases and controls (p < 1×10”^6^) as well as censored on minor allele frequency (MAF > 0.01). In total 8,018,013 imputed dosages and best guess genotypes were used for analysis.

Two cohorts of unrelated subjects with homogenous genetic ancestry were created for GWAS analysis (GWAS cohort) and GREML SNP-heritability analysis (GCTA cohort). Genetic ancestry was characterized using principal components (PC) analysis implemented in smartPCA^104,105^. We performed two iterations of censoring, removing subjects outlying from joint distribution of the first 10 PCs defined in the subset of iPSYCH with four grandparents recorded in the Danish civil register as born in Denmark, re-computing PCs on the remaining subjects and censoring again according to the same criteria. Censored individuals were aggregated into a replication cohort. For the GWAS cohort kinship was estimated using KING^106^ and individuals were censored to ensure no pair had closer than third degree kinship. For the GCTA cohort, kinship was more strictly filtered such that no pair had GCTA-based estimate greater than 0.034, the absolute value of the minimum estimated kinship. When possible cases were retained and the control relative was censored. All subject genotypes were checked for abnormal sample heterozygosity, high levels of missing genotypes (> 1%), sex concordance and inconsistencies among duplicate samples. In total 65,534 subjects were retained in the GWAS cohort, 43,311 in the GCTA cohort and 7,163 in the Replication cohort. A detailed genotype and subject quality control protocol is provided as a Supplementary Note.

### SNP heritability and genetic correlations

SNP-heritability and genetic correlations were estimated in the GCTA cohort with the GREML approach available in GCTA^107–109^. Age, gender and ten principal components were included as fixed effects covariates. Single indication estimates of SNP heritability were estimated using the same random population controls and each individual case group. Estimates were converted to the liability scale^109^ according to estimates of population prevalence made in the random population portion of the GCTA cohort (Supplementary Table 2). Estimation of genetic correlation between indications was performed using bivariate GREML^108,110^. For each pair, subjects with both indications were excluded and controls were randomly and evenly split. Splitting and estimation were repeated 5 times for each pair and the median values were retained.

Published GCTA SNP heritability estimates for ADHD, AFF, ASD, BIP, SCZ were taken from Lee et al^15^. GREML estimates of SNP-for ANO and XDX were unavailable. GWAS statistics for eXDX^34^, eADHD^39^, eAFF^41^, eANO^42^, eASD^40^, and eSCZ^44^ were downloaded from the Psychiatric Genomics Consortium (PGC) repository (http://www.med.unc.edu/pgc/results-and-downloads). eBIP^43^ statistics were downloaded from the NHGRI-EBI GWAS catalog^47^. Linkage disequilibrium score regression (LDSC)^111^ was used to estimate SNP heritability for these published studies and for each single iPSYCH indication GWAS. Reference LD scores and protocol were provided by the authors (https://github.com/bulik/ldsc/wiki). Genetic correlations between iPSYCH indications and published GWAS were estimated with LDSC^16^ using the authors' protocols. To facilitate comparisons a typical population prevalence was used for each liability scale transformation (XDX=0.35, ADHD=0.05, AFF=0.15, ANO=0.01, ASD=0.01, BIP=0.01, SCZ=0.01)^42,45^, including re-scaling the iPSYCH GREML estimates.

### Association Testing

GWAS were performed using imputed additive genotype dosages and logistic regression implemented in plink version 2^112–113^. The XDX GWAS included all subjects in the GWAS cohort (46,008 cases, 19,526 controls). Age, gender and ten principal components were included as fixed effects covariates. Stratified false discovery rates^114^ (sFDR) were estimated according to Story's q-value^115^ and computed independently for common (minor allele frequency, MAF, ≥ 0.05) and uncommon SNPs (0.01 < MAF < 0.05). The suggestive SNP threshold of sFDR q-value < 0.05 corresponds to a p-value less than 1.02 × 10^−5^ for common SNPs and less than 4.71 × 10^−7^ for uncommon SNPs. Secondary GWAS including or excluding a single indication were performed via the same protocols and sharing the same 19,526 controls. These analyses were performed only to provide context for the XDX results. For the internal replication cohort (7,163 individuals, 4,481 cases), association tests used best guess genotypes and linear mixed models implemented in GCTA^108^ accounting for relatedness and heterogeneity in genetic background with genome-wide estimates of empirical kinship. Gender and age were included as fixed effects covariates.

### Fine Mapping

Region based loci associated with independent index SNPs were defined and refined iteratively. The most significant SNP was selected and plink2^112,113^ was used to estimate pairwise r^2^ LD between the index SNP and all SNPs within 5 megabases. All SNPs with r^2^ LD > 0.1 with the index SNP were considered index-associated SNPs. Locus bounds were determined by the physical positions of the furthest index-associated SNP up and downstream. The process was repeated until all suggestive SNPs were labeled as index or index-associated SNPs. Loci with overlapping index-associated SNP sets were merged. Conditional analysis was performed within each locus, including the imputed dosage for the most significant SNP as a covariate and re-computing within locus association statistics. Secondary suggestive SNPs were retained as independent index SNPs and the process was repeated until no SNPs within the locus showed suggestive association. Credible SNPs were defined for each locus using FineMap^62^ with default parameters. For each locus, FineMap input SNPs had LD r^2^ > 0.6 with the index SNP and an association p-value less than 0.001. Using the per SNP posterior probabilities and Bayes factors provided by FineMap, we define credible sets as the minimum collection of SNPs needed to achieve 95% posterior inclusion probability, supplemented with the index SNP and any individual SNPs with log_10_(Bayes Factors) greater than 2.

### External Statistical Support

The 13 November 2017 NHGRI GWAS catalog^47^ contains 58,215 autosomal single SNP associations with positions that could be mapped to the hg19 reference aggregated from 3,893 GWAS described in 2,278 publications. 539 GWAS were labeled as “Neurological Disease,” a broad category including psychiatric outcomes, 3,354 were not. Catalog SNPs were connected to XDX loci according to an objective hierarchy. First, catalog SNPs were checked if they were an index SNP. If not, they were checked if they were a credible SNP. Remaining catalog SNPs were checked for r^2^ LD > 0.6 with a credible SNP using iPSYCH genotypes. If catalog SNPs were not present in the iPSYCH genotypes, we checked for an LD connection in the 1000 genomes project phase 3^103^. Effect directions were aligned to the same strand by allele codes for unambiguous SNPs (A-T/C-G) and by frequency when the minor allele frequency was less than 0.40 for strand ambiguous SNPs (A-T/T-A, C-G/G-C). Connections to strand and minor allele ambiguous SNPs are noted by unsigned p-values in Supplementary Table 17. Enrichment for connections to neurological GWAS was tested with a binomial proportion test, although this test may not be optimally specified due to overlap among cataloged studies

To connect each locus to the full results from the seven external GWAS described previously, we followed a similar protocol. Priority was given to index SNPs genotyped in both studies, then credible SNPs, then the strongest proxy-credible LD pair with r^2^ at least 0.6 in the iPSYCH data. Only strand unambiguous SNPs were considered (A-T/C-G). For concordance tests, we considered effects of the proxy SNP in both studies to ensure we used statistics for the same variant in the case where an index or credible SNP was not directly present in the external GWAS. Concordance was estimated via a linear model predicting the external z-score from the XDX z-score, including an intercept. The same concordance test was used for results from the internal replication association tests.

### LDSC-SEG

LDSC regression for specifically expressed genes (LDSC-SEG)^61^ tests for enrichments in per SNP heritability among sets of SNPs defined for plausibly tissue preferential biological signatures while controlling for tissue general effects. Three sets of pre-computed annotation files (LD scores) are provided by the LDSC-SEG authors (GTEx, DEPICT and RoadMap). The GTEx LD-scores represent 53 variant annotations capturing preferentially expressed genes defined from human tissue RNA sequencing^116^. The DEPICT LD-scores represent 152 variant annotations capturing tissue preferential gene sets defined from an amalgamation of human, mouse, and rat microarray experiments^117^. The Roadmap LD-scores represent 396 variant annotations constructed from data produced by the Roadmap Epigenetics project^118^, namely the narrow peaks defined by Roadmap for DNase hypersensitivity, H3K27ac, H3K4me3, H3K4me1, H3K9ac, and/or H3K36me3 chromatin marks in 88 cell types or primary tissues. Analytic protocols were as provided by the LDSC-SEG developers documentation (https://github.com/bulik/ldsc/wiki/Cell-type-specific-analyses). P-values less than 8.32×10^−5^ were declared significant to correct for testing 601 sets of LD scores, and suggestive significance was determined by Story's q-value^115^ (FDR < 0.05 which corresponds to p < 8.27 × 10^−4^). Replication LDSC-SEG partitioning used the eSCZ^44^ statistics. Internal replication enrichments used results from seven secondary GWAS, each one censoring all patients with a different indication.

### Identifying Candidate Genes

For each locus, candidate genes were identified by functional connections between credible SNPs and plausible targets according to the union of three selection criteria as described previously^63^. First, genes containing credible SNPs that cause missense variation or nonsense mediated decay were selected (133 credible SNPs implicating 13 candidate genes). Second, genes with credible SNPs located in the promoter regions were selected (2 kilobases upstream from the transcription start site; 15 credible SNPs implicating 12 candidate genes). Finally, unannotated SNPs were assigned to genes based on three-dimensional chromatin contacts defined by an interaction map for fetal brain^63^. Genes contacting regions containing credible SNPs were selected (252 implicating 262 candidate genes). In total, 400 credible SNPs were assigned to 281 candidate genes.

### Candidate Gene Enrichment Tests

Gene ontology enrichments were performed using GOElite^119^ and ontologies from ENSMart^120^ v77 against a background of all autosomal protein coding genes. Developmental expression trajectories for candidate genes were plotted using a published transcriptome atlas constructed from post mortem cortices^121^. Expression values were log-transformed and centered using the mean expression values for all brain expressed genes. Mean expression values for the 281 candidate genes were plotted across prenatal (6–37 weeks post-conception) and postnatal (4 months to 42 years) developmental stages. Developing neural cell-type enrichments were estimated using expression profiles of single-cells taken from fetal cortical laminae^66^. Cell-type specific genes were selected according to a significant (FDR < 0.05) Pearson correlation between the gene and an idealized cluster marker for each cell-type, following the approach described in the data generation report^66^. Candidate gene enrichment for each set of specifically expressed genes was estimated by logistic regression and adjusting for gene length.

## Acknowledgements

The iPSYCH Initiative is funded by the Lundbeck Foundation (grant numbers R102-A9118 and R155–2014–1724), the Mental Health Services Capital Region of Denmark, University of Copenhagen, Aarhus University and The university hospital in Aarhus. Genotyping of iPSYCH samples was supported by grants from the Lundbeck Foundation, the Stanley Foundation, the Simons Foundation (SFARI 311789), and NIMH (5U01MH094432–02). The IPSYCH Initiative utilize the Danish National Biobank resource that is supported by the Novo Nordisk Foundation. IPSYCH data was stored and analyzed at the Computerome HPC facility (http://www.computerome.dtu.dk/) and authors are grateful for continuous support from the HPC team led by Dr. AN Syed. HW was supported by NIH grant K99MH113823. WKT is in part supported by NIH grant R01GM104400.

## Author Contributions

D. G. and T.W. conceived of and supervised the study. A.J.S., H.W., D.G. and T.W. designed the analysis plan. A.J.S., V.A., A.B and W.K.T. prepared the data. A.J.S. performed the GWAS, (partitioned) SNP-(co)heritability, fine-mapping and replication analyses. H.W. performed the candidate gene identification and enrichment analyses. R.N. and M.G. provided interpretive support. O.D. contributed imputation software and protocols. D.M.H, M.B-H., J.B-G., M.G.P., E. A., C.B.P., B.M.N., M.J.D., M.N., O.M., A.D.B., P.B.M and T.W. designed, implemented and/or oversaw the collection and generation of the iPSYCH data. A.J.S., H.W., D.G. and T.W. wrote the manuscript. All authors discussed the results and contributed to the revision of the manuscript.

## Supplementary Table Legends

Supplementary Table 1. Indication Descriptive Statistics. Here we report the number of individuals from each indication group for each iPSYCH cohort used in our analyses. We show the breakdown by gender and summarize the distribution of age for each as well. *Age as of 1/1/2014, through when diagnoses are complete. SD, Standard Deviation.

Supplementary Table 2. Indication Prevalence Estimates. Here we report the population prevalence estimates for each indication group. We estimate the population prevalence as the proportion of indication positive individuals in portion of the cohort that was ascertained as a true random population sample.

Supplementary Table 3. iPSYCH Indication SNP-heritability. Here we provide the statistics that were used to create Figure 1A. SNP-heritability was estimate via GREML using all individuals with each indication. Liability transformations used the population prevalence estimates from the GCTA cohort as shown in Supplementary Table 2.

Supplementary Table 4. SNP-based genetic correlations from multiple sources and methods. Here we provide the statistics that were used to create Figure 1B. GREML SNP-heritability was estimated using the GCTA cohort and all subjects with each indication. External estimates were taken from Lee et al^45^. iPSYCH LDSC SNP-heritability estimates were computed from GWAS of all subjects with each individual indication following the same protocol as the XDX GWAS. External LDSC estimates were computed using eGWAS statistics described previously. LD-scores and analysis protocol were as provided by the authors. Liability transformations were according to commonly stated population prevalence estimates, including re-scaling iPSYCH GREML estimates from Figure 1A and Supplementary Table 3.

Supplementary Table 5. iPSYCH-eGWAS genetic correlations. Here we provide the statistics that were used to create Figure 1C. We used LDSC to estimate the genetic correlation between iPSYCH indications and external studies of the same disorder. For iPSYCH, we used statistics from GWAS performed via the same protocol as the XDX but limiting cases to all subjects with a given indication.

Supplementary Table 6. Median iPSYCH across indication genetic correlations. Here we provide the statistics that were used to create the blue bars in Figure 1D. We used bivariate GREML to estimate the genetic correlation between iPSYCH indications, for each pair censoring subjects with both diagnoses and randomly splitting controls. These statistics show only the median values for each repeated pair.

Supplementary Table 7. All iPSYCH across indication genetic correlations. We used bivariate GREMLto estimate the genetic correlation between iPSYCH indications, for each pair censoring subjects with both diagnoses and randomly splitting controls. These statistics show all estimations.

Supplementary Table 8. All eGWAS across disorder genetic correlations. Here we provide the statistics that were used to create the grey bars in Figure 1D. We used bivariate LDSC to estimate the genetic correlation between each eGWAS, using the LD-scores and protocol made available by the LDSC authors.

Supplementary Table 9. XDX GWAS association statistics for the top 50 loci. Here we provide the index SNP association statistics and gene connections for each of the top 50 loci. All loci are primary associations, except for locus 13 which was identified as a secondary suggestive locus within locus 3.

Supplementary Table 10. Single indication (all subjects) GWAS association statistics for the top 50 loci. Here we provide the index SNP association statistics for each of the top 50 loci from association tests using all patients with each single indication as cases.

Supplementary Table 11. Single indication (exclusive subjects) GWAS association statistics for the top 50 loci. Here we provide the index SNP association statistics for each of the top 50 loci from association tests using only patients with a single indication as cases.

Supplementary Table 12. Statistics summarizing all connections between the top 50 XDX loci and the GWAS catalog. Here we describe the associations summarized in the GWAS catalog and their connection to each of our top 50 loci.

Supplementary Table 13. Statistics summarizing all connections between the top 50 XDX loci and the eGWAS statistics. Here we describe the associations summarized in each of the eGWAS described previously and their connection to each of our top 50 loci.

Supplementary Table 14. Replication Statistics for Index SNPs from the top 50 XDX loci. Here we present the associations statistics from the linear mixed model replication association in the small, independent and ancestry-diverse iPSYCH replication cohort.

Supplementary Table 15. LDSC-SEG Statistics for the GTEx LD-scores. Here we present the enrichment statistics from the LDSC-SEG analysis of the XDX GWAS using the GTEx LD-scores. The statistics were used to make Supplementary Figure 11.

Supplementary Table 16. LDSC-SEG Statistics for the DEPICT LD-scores. Here we present the enrichment statistics from the LDSC-SEG analysis of the XDX GWAS using the DEPICT LD-scores. The statistics were used to make Supplementary Figure 12.

Supplementary Table 17. LDSC-SEG statistics for the CNS RoadMap LD-scores. Here we present the enrichment statistics from the LDSC-SEG analysis of the XDX GWAS using the CNS RoadMap LD-scores. The statistics were used to make Figure 4A.

Supplementary Table 18. LDSC-SEG statistics for the all RoadMap LD-scores. Here we present the enrichment statistics from the LDSC-SEG analysis of the XDX GWAS using all RoadMap LD-scores. The statistics were used to make Supplementary Figure 13.

Supplementary Table 19. LDSC-SEG statistics for the CNS RoadMap LD-scores in the eSCZ GWAS. Here we present the enrichment statistics from the LDSC-SEG analysis of the eSCZ GWAS using the CNS RoadMap LD-scores. The statistics were used to make Figure 4B.

Supplementary Table 20. LDSC-SEG statistics for the CNS RoadMap LD-scores in the leave-one-out XDX GWAS. Here we present the enrichment statistics from the LDSC-SEG analysis for the XDX phenotype, iteratively dropping each individual indication. The statistics were used to make Figure 4C.

Supplementary Table 21. Association statistics and gene connections for credible SNPs. Here we present the XDX GWAS association statistics, FineMap results and functional gene connections for each of 627 credible SNPs.

Supplementary Table 22. Candidate gene functions. Here we aggregate Brief refSeq^80^ and UniProt^81^ descriptions for all candidate protein coding genes were aggregated from UCSC genome browser databases^82,83^.

## References

1 Robins, E. & Guze, S. B. Establishment of diagnostic validity in psychiatric illness: its application to schizophrenia. Am J Psychiatry 126, 983–987, doi:10.1176/ajp.126.7.983 (1970).

2 Kendell, R. & Jablensky, A. Distinguishing between the validity and utility of psychiatric diagnoses. Am J Psychiatry 160, 4–12, doi:10.1176/appi.ajp.160.1.4 (2003).

3 Krystal, J. H., & State, M. W. Psychiatric disorders: diagnosis to therapy. Cell 157, 201–214, doi:10.1016/j.cell.2014.02.042 (2014).

4 O'Donovan, M. C. & Owen, M. J. The implications of the shared genetics of psychiatric disorders. Nature medicine 22,1214–1219, doi:10.1038/nm.4196 (2016).

5 Doherty, J. L. & Owen, M. J. Genomic insights into the overlap between psychiatric disorders: implications for research and clinical practice. Genome Med 6, 29, doi:10.1186/gm546 (2014).

6 Henin, A. et al. Childhood antecedent disorders to bipolar disorder in adults: a controlled study. J Affect Disord 99, 51–57, doi:10.1016/j.jad.2006.09.001 (2007).

7 Widiger, T. A., & Sankis, L. M. Adult psychopathology: issues and controversies. Annu Rev Psychol 51, 377–404, doi:10.1146/annurev.psych.51.1.377 (2000).

8 Bulik, C. M., Prescott, C. A. & Kendler, K. S. Features of childhood sexual abuse and the development of psychiatric and substance use disorders. Br J Psychiatry 179, 444–449 (2001).

9 Brown, G. W., Harris, T. O. & Eales, M. J. Social factors and comorbidity of depressive and anxiety disorders. Br J Psychiatry Suppl, 50–57 (1996).

10 Gorman, J. M. & Kent, J. M. SSRIs and SNRIs: broad spectrum of efficacy beyond major depression. J Clin Psychiatry 60 Suppl 4, 33–38; discussion 39 (1999).

11 Polderman, T. J. et al. Meta-analysis of the heritability of human traits based on fifty years of twin studies. Nat Genet 47, 702–709, doi:10.1038/ng.3285 (2015).

12 Lichtenstein, P. et al. Common genetic determinants of schizophrenia and bipolar disorder in Swedish families: a population-based study. Lancet 373, 234–239, doi:10.1016/S0140-6736(09)60072-6 (2009).

13 Sullivan, P. F. et al. Family history of schizophrenia and bipolar disorder as risk factors for autism. Archives of general psychiatry 69,1099–1103, doi:10.1001/archgenpsychiatry.2012.730 (2012).

14 Larsson, H. et al. Risk of bipolar disorder and schizophrenia in relatives of people with attention-deficit hyperactivity disorder. Br J Psychiatry 203,103–106, doi:10.1192/bjp.bp. 112.120808 (2013).

15 Cross-Disorder Group of the Psychiatric Genomics, C. et al. Genetic relationship between five psychiatric disorders estimated from genome-wide SNPs. Nat Genet 45, 984–994, doi:10.1038/ng.2711 (2013).

16 Bulik-Sullivan, B. et al. An atlas of genetic correlations across human diseases and traits. Nat Genet 47, 1236–1241, doi:10.1038/ng.3406 (2015).

17 International Schizophrenia, C. et al. Common polygenic variation contributes to risk of schizophrenia and bipolar disorder. Nature 460, 748–752, doi:10.1038/nature08185 (2009).

18 Wray, N. R. & Sullivan, P. F. Genome-wide association analyses identify 44 risk variants and refine the genetic architecture of major depression. bioRxiv, doi:10.1101/167577 (2017).

19 Hamshere, M. L. et al. Shared polygenic contribution between childhood attention-deficit hyperactivity disorder and adult schizophrenia. Br J Psychiatry 203,107–111, doi:10.1192/bjp.bp. 112.117432 (2013).

20 Demontis, D. et al. Discovery Of The First Genome-Wide Significant Risk Loci For ADHD. bioRxiv, doi:10.1101/145581 (2017).

21 Grove, J. et al. Common risk variants identified in autism spectrum disorder. bioRxiv, doi:10.1101/224774 (2017).

22 Kirov, G. et al. The penetrance of copy number variations for schizophrenia and developmental delay. Biol Psychiatry 75, 378–385, doi:10.1016/j.biopsych.2013.07.022 (2014).

23 Green Snyder, L. et al. Autism Spectrum Disorder, Developmental and Psychiatric Features in 16p11.2 Duplication. J Autism Dev Disord 46, 2734–2748, doi:10.1007/s10803-016-2807-4 (2016).

24 Schneider, M. et al. Psychiatric disorders from childhood to adulthood in 22q11.2 deletion syndrome: results from the International Consortium on Brain and Behavior in 22q11.2 Deletion Syndrome. Am J Psychiatry 171, 627–639, doi:10.1176/appi.ajp.2013.13070864 (2014).

25 Pak, C. et al. Human Neuropsychiatric Disease Modeling using Conditional Deletion Reveals Synaptic Transmission Defects Caused by Heterozygous Mutations in NRXN1. Cell Stem Cell 17, 316–328, doi:10.1016/j.stem.2015.07.017 (2015).

26 Rujescu, D. et al. Disruption of the neurexin 1 gene is associated with schizophrenia. Hum Mol Genet 18, 988–996, doi:10.1093/hmg/ddn351 (2009).

27 Ching, M. S. et al. Deletions of NRXN1 (neurexin-1) predispose to a wide spectrum of developmental disorders. Am J Med Genet B Neuropsychiatr Genet 153B, 937–947, doi:10.1002/ajmg.b.31063 (2010).

28 Ferreira, M. A. et al. Collaborative genome-wide association analysis supports a role for ANK3 and CACNA1C in bipolar disorder. Nat Genet 40,1056–1058, doi:10.1038/ng.209 (2008).

29 Schizophrenia Psychiatric Genome-Wide Association Study, C. Genome-wide association study identifies five new schizophrenia loci. Nat Genet 43, 969–976, doi:10.1038/ng.940 (2011).

30 O'Donovan, M. C. et al. Identification of loci associated with schizophrenia by genome-wide association and follow-up. Nat Genet 40,1053–1055, doi:10.1038/ng.201 (2008).

31 Williams, H. J. et al. Fine mapping of ZNF804A and genome-wide significant evidence for its involvement in schizophrenia and bipolar disorder. Mol Psychiatry 16, 429–441, doi:10.1038/mp.2010.36 (2011).

32 Cichon, S. et al. Genome-wide association study identifies genetic variation in neurocan as a susceptibility factor for bipolar disorder. Am J Hum Genet 88, 372–381, doi:10.1016/j.ajhg.2011.01.017 (2011).

33 Muhleisen, T. W. et al. Association between schizophrenia and common variation in neurocan (NCAN), a genetic risk factor for bipolar disorder. Schizophrenia research 138, 69–73, doi:10.1016/j.schres.2012.03.007 (2012).

34 Cross-Disorder Group of the Psychiatric Genomics, C. Identification of risk loci with shared effects on five major psychiatric disorders: a genome-wide analysis. Lancet 381, 1371–1379, doi:10.1016/S0140-6736(12)62129-1 (2013).

35 Green, E. K. et al. The bipolar disorder risk allele at CACNA1C also confers risk of recurrent major depression and of schizophrenia. Mol Psychiatry 15,1016–1022, doi:10.1038/m p.2009.49 (2010).

36 Psychiatric, G. C. B. D. W. G. Large-scale genome-wide association analysis of bipolar disorder identifies a new susceptibility locus near ODZ4. Nat Genet 43, 977–983, doi: 10.1038/ng.943 (2011).

37 Gandal, M. J. et al. Shared molecular neuropathology across major psychiatric disorders parallels polygenic overlap. Science (In Press).

38 Pedersen, C. B. et al. The iPSYCH2012 case-cohort sample: new directions for unravelling genetic and environmental architectures of severe mental disorders. Mol Psychiatry, doi :10.1038/m p.2017.196 (2017).

39 Neale, B. M. et al. Meta-analysis of genome-wide association studies of attention-deficit/hyperactivity disorder. J Am Acad Child Adolesc Psychiatry 49, 884–897, doi:10.1016/j.jaac.2010.06.008 (2010).

40 Autism Spectrum Disorders Working Group of The Psychiatric Genomics, C. Meta-analysis of GWAS of over 16,000 individuals with autism spectrum disorder highlights a novel locus at 10q24.32 and a significant overlap with schizophrenia. Mol Autism 8, 21, doi:10.1186/s13229-017-0137-9 (2017).

41 Major Depressive Disorder Working Group of the Psychiatric, G. C. et al. A mega-analysis of genome-wide association studies for major depressive disorder. Mol Psychiatry 18, 497–511, doi: 10.1038/m p.2012.21 (2013).

42 Duncan, L. et al. Significant Locus and Metabolic Genetic Correlations Revealed in Genome-Wide Association Study of Anorexia Nervosa. Am J Psychiatry 174, 850–858, doi:10.1176/appi.ajp.2017.16121402 (2017).

43 Hou, L. et al. Genome-wide association study of 40,000 individuals identifies two novel loci associated with bipolar disorder. Hum Mol Genet 25, 3383–3394, doi:10.1093/hmg/ddw181 (2016).

44 Schizophrenia Working Group of the Psychiatric Genomics, C. Biological insights from 108 schizophrenia-associated genetic loci. Nature 511, 421–427, doi:10.1038/nature13595 (2014).

45 Cross-Disorder Group of the Psychiatric Genomics Consortium et al. Genetic relationship between five psychiatric disorders estimated from genome-wide SNPs. Nat Genet 45, 984–994, doi:10.1038/ng.2711 (2013).

46 Devlin, B. & Roeder, K. Genomic control for association studies. Biometrics 55, 997–1004 (1999).

47 MacArthur, J. et al. The new NHGRI-EBI Catalog of published genome-wide association studies (GWAS Catalog). Nucleic Acids Res 45, D896–D901, doi:10.1093/nar/gkw1133 (2017).

48 Guo, W. et al. Polygenic risk score and heritability estimates reveals a genetic relationship between ASD and OCD. Eur Neuropsychopharmacol 27, 657–666, doi:10.1016/j.euroneuro.2017.03.011 (2017).

49 Goes, F. S. et al. Genome-wide association study of schizophrenia in Ashkenazi Jews. Am J Med Genet B Neuropsychiatr Genet 168, 649–659, doi:10.1002/ajmg.b.32349 (2015).

50 Hek, K. et al. A genome-wide association study of depressive symptoms. Biol Psychiatry 73, 667–678, doi:10.1016/j.biopsych.2012.09.033 (2013).

51 van't Hof, F. N. et al. Shared Genetic Risk Factors of Intracranial, Abdominal, and Thoracic Aneurysms. J Am Heart Assoc 5, doi:10.1161/JAHA.115.002603 (2016).

52 Bilguvar, K. et al. Susceptibility loci for intracranial aneurysm in European and Japanese populations. Nat Genet 40,1472–1477, doi:10.1038/ng.240 (2008).

53 Wakil, S. M. et al. A common variant association study reveals novel susceptibility loci for low HDL-cholesterol levels in ethnic Arabs. Clin Genet 90, 518–525, doi:10.1111/cge. 12761 (2016).

54 Perry, J. R. et al. Parent-of-origin-specific allelic associations among 106 genomic loci for age at menarche. Nature 514, 92–97, doi:10.1038/nature13545 (2014).

55 Astle, W. J. et al. The Allelic Landscape of Human Blood Cell Trait Variation and Links to Common Complex Disease. Cell 167,1415–1429 e1419, doi:10.1016/j.cell.2016.10.042 (2016).

56 Keene, K. L. et al. Genetic Associations with Plasma B12, B6, and Folate Levels in an Ischemic Stroke Population from the Vitamin Intervention for Stroke Prevention (VISP) Trial. Front Public Health 2,112, doi:10.3389/fpubh.2014.00112 (2014).

57 Hazra, A. et al. Genome-wide significant predictors of metabolites in the one-carbon metabolism pathway. Hum Mol Genet 18, 4677–4687, doi:10.1093/hmg/ddp428 (2009).

58 van Meurs, J. B. et al. Common genetic loci influencing plasma homocysteine concentrations and their effect on risk of coronary artery disease. Am J Clin Nutr 98, 668–676, doi:10.3945/ajcn.112.044545 (2013).

59 Jin, Y. et al. Genome-wide association studies of autoimmune vitiligo identify 23 new risk loci and highlight key pathways and regulatory variants. Nat Genet 48,1418–1424, doi:10.1038/ng.3680 (2016).

60 Hyde, C. L. et al. Identification of 15 genetic loci associated with risk of major depression in individuals of European descent. Nat Genet 48,1031–1036, doi:10.1038/ng.3623 (2016).

61 Finucane, H. et al. Heritability enrichment of specifically expressed genes identifies disease-relevant tissues and cell types. bioRxiv (2017).

62 Benner, C. et al. FINEMAP: efficient variable selection using summary data from genome-wide association studies. Bioinformatics 32,1493–1501, doi:10.1093/bioinformatics/btw018 (2016).

63 Won, H. et al. Chromosome conformation elucidates regulatory relationships in developing human brain. Nature 538, 523–527, doi:10.1038/nature19847 (2016).

64 Parikshak, N. N. et al. Integrative functional genomic analyses implicate specific molecular pathways and circuits in autism. Cell 155,1008–1021, doi:10.1016/j.cell.2013.10.031 (2013).

65 Network & Pathway Analysis Subgroup of Psychiatric Genomics, C. Psychiatric genome-wide association study analyses implicate neuronal, immune and histone pathways. Nat Neurosci 18, 199–209, doi:10.1038/nn.3922 (2015).

66 Pollen, A. A. et al. Molecular identity of human outer radial glia during cortical development. Cell 163, 55–67, doi:10.1016/j.cell.2015.09.004 (2015).

67 Bender, A. T., & Beavo, J. A. Cyclic nucleotide phosphodiesterases: molecular regulation to clinical use. Pharmacol Rev 58, 488–520, doi:10.1124/pr.58.3.5 (2006).

68 Wishart, D. S. et al. DrugBank 5.0: a major update to the DrugBank database for 2018. Nucleic Acids Res, doi:10.1093/nar/gkx1037 (2017).

69 Han, Q. J. et al. IPP5 inhibits neurite growth in primary sensory neurons by maintaining TGF-beta/Smad signaling. J Cell Sci 126, 542–553, doi:10.1242/jcs.H4280 (2013).

70 Munton, R. P., Vizi, S. & Mansuy, I. M. The role of protein phosphatase-1 in the modulation of synaptic and structural plasticity. FEBS Lett 567,121–128, doi:10.1016/j.febslet.2004.03.121 (2004).

71 Kosmicki, J. A. et al. Refining the role of de novo protein-truncating variants in neurodevelopmental disorders by using population reference samples. Nat Genet 49, 504–510, doi:10.1038/ng.3789 (2017).

72 Sanders, S. J. et al. Insights into Autism Spectrum Disorder Genomic Architecture and Biology from 71 Risk Loci. Neuron 87, 1215–1233, doi:10.1016/j.neuron.2015.09.016 (2015).

73 Jang, S. et al. Synaptic adhesion molecule IgSFll regulates synaptic transmission and plasticity. Nat Neurosci 19, 84–93, doi:10.1038/nn.4176 (2016).

74 Hermey, G. The Vps10p-domain receptor family. Cell Mol Life Sci 66, 2677–2689, doi:10.1007/s00018-009-0043-1 (2009).

75 Breiderhoff, T. et al. Sortilin-related receptor SORCS3 is a postsynaptic modulator of synaptic depression and fear extinction. PLoS One 8, e75006, doi:10.1371/journal.pone.0075006 (2013).

76 O'Sullivan, M. L., Martini, F., von Daake, S., Comoletti, D. & Ghosh, A. LPHN3, a presynaptic adhesion-GPCR implicated in ADHD, regulates the strength of neocortical layer 2/3 synaptic input to layer 5. Neural Dev 9, 7, doi:10.1186/1749-8104-9-7 (2014).

77 Tomioka, N. H. et al. Elfn1 recruits presynaptic mGluR7 in trans and its loss results in seizures. Nat Commun 5, 4501, doi:10.1038/ncomms5501 (2014).

78 Christensen, E. I., & Birn, H. Megalin and cubilin: multifunctional endocytic receptors. Nat Rev Mol Cell Biol 3, 256–266, doi:10.1038/nrm778 (2002).

79 Grasbeck, R. Imerslund-Grasbeck syndrome (selective vitamin B(12) malabsorption with proteinuria). OrphanetJ Rare Dis 1,17, doi:10.1186/1750-1172-1-17 (2006).

80 Pruitt, K. D. et al. RefSeq: an update on mammalian reference sequences. Nucleic Acids Res 42, D756–763, doi:10.1093/nar/gkt1114 (2014).

81 UniProt, C. Reorganizing the protein space at the Universal Protein Resource (UniProt). Nucleic Acids Res 40, D71–75, doi:10.1093/nar/gkr981 (2012).

82 Tyner, C. et al. The UCSC Genome Browser database: 2017 update. Nucleic Acids Res 45, D626–D634, doi:10.1093/nar/gkw1134 (2017).

83 Karolchik, D. et al. The UCSC Table Browser data retrieval tool. Nucleic Acids Res 32, D493–496, doi:10.1093/nar/gkh103 (2004).

84 Wray, N. R., Lee, S. H. & Kendler, K. S. Impact of diagnostic misclassification on estimation of genetic correlations using genome-wide genotypes. EurJ Hum Genet 20, 668–674, doi:10.1038/ejhg.2011.257 (2012).

85 Sudlow, C. et al. UK biobank: an open access resource for identifying the causes of a wide range of complex diseases of middle and old age. PLoS medicine 12, e1001779, doi:10.1371/journal.pmed.1001779 (2015).

86 Wang, K., Gaitsch, H., Poon, H., Cox, N. J. & Rzhetsky, A. Classification of common human diseases derived from shared genetic and environmental determinants. Nat Genet 49, 1319–1325, doi:10.1038/ng.3931 (2017).

87 Polubriaginof, F. et al. Estimate of disease heritability using 7.4 million familial relationships inferred from electronic health records. bioRxiv, doi:10.1101/066068 (2017).

88 Mortensen, P. B. et al. Effects of family history and place and season of birth on the risk of schizophrenia. N Engl J Med 340, 603–608, doi:10.1056/NEJM199902253400803 (1999).

89 Giovanoli, S. et al. Stress in puberty unmasks latent neuropathological consequences of prenatal immune activation in mice. Science 339,1095–1099, doi:10.1126/science. 1228261 (2013).

90 Lindenbaum, J. et al. Neuropsychiatric disorders caused by cobalamin deficiency in the absence of anemia or macrocytosis. N Engl J Med 318,1720–1728, doi: 10.1056/NEJM198806303182604 (1988).

91 Briani, C. et al. Cobalamin deficiency: clinical picture and radiological findings. Nutrients 5, 4521–4539, doi:10.3390/nu5114521 (2013).

92 Dommisse, J. Subtle vitamin-B12 deficiency and psychiatry: a largely unnoticed but devastating relationship? Med Hypotheses 34,131–140 (1991).

93 Mitchell, E. S., Conus, N. & Kaput, J. B vitamin polymorphisms and behavior: evidence of associations with neurodevelopment, depression, schizophrenia, bipolar disorder and cognitive decline. Neurosci Biobehav Rev 47, 307–320, doi:10.1016/j.neubiorev.2014.08.006 (2014).

94 Pedersen, C. B. The Danish Civil Registration System. Scand J Public Health 39, 22–25, doi:10.1177/1403494810387965 (2011).

95 Lynge, E., Sandegaard, J. L. & Rebolj, M. The Danish National Patient Register. ScandJ Public Health 39, 30–33, doi:10.1177/1403494811401482 (2011).

96 Mors, O., Perto, G. P. & Mortensen, P. B. The Danish Psychiatric Central Research Register. Scand J Public Health 39, 54–57, doi:10.1177/1403494810395825 (2011).

97 Norgaard-Pedersen, B. & Hougaard, D. M. Storage policies and use of the Danish Newborn Screening Biobank. J Inherit Metab Dis 30, 530–536, doi:10.1007/s10545-007-0631-x (2007).

98 Pedersen, C. B. et al. A comprehensive nationwide study of the incidence rate and lifetime risk for treated mental disorders. JAMA Psychiatry 71, 573–581, doi:10.1001/jamapsychiatry.2014.16 (2014).

99 World Health Organization. The ICD-10 Classification of Mental and Behavioural Disorders. Diagnostic criteria for research. (Geneva).

100 World Health Organization. Klassifikation af sygdomme; Udvidet dansk-latinsk udgave af verdenssundhedsorganisationens internationale klassifikation af sygdomme. 8 revision, 1965 [Classification of diseases: Extended Danish-Latin version of the World Health Organization International Classification of Diseases, 8th revision, 1965]. (Danish National Board of Health, Copenhagen, 1971).

101 O'Connell, J. et al. Haplotype estimation for biobank-scale data sets. Nat Genet 48, 817–820, doi:10.1038/ng.3583 (2016).

102 Howie, B. N., Donnelly, P. & Marchini, J. A flexible and accurate genotype imputation method for the next generation of genome-wide association studies. PLoS Genet 5, e1000529, doi:10.1371/journal.pgen.1000529 (2009).

103 1000 Genomes Project Consortium et al. A global reference for human genetic variation. Nature 526, 68–74, doi:10.1038/nature15393 (2015).

104 Price, A. L. et al. Principal components analysis corrects for stratification in genome-wide association studies. Nat Genet 38, 904–909, doi:10.1038/ng1847 (2006).

105 Patterson, N., Price, A. L. & Reich, D. Population structure and eigenanalysis. PLoS Genet 2, e190, doi:10.1371/journal.pgen.0020190 (2006).

106 Manichaikul, A. et al. Robust relationship inference in genome-wide association studies. Bioinformatics 26, 2867–2873, doi:10.1093/bioinformatics/btq559 (2010).

107 Yang, J. et al. Common SNPs explain a large proportion of the heritability for human height. Nat Genet 42, 565–569, doi:10.1038/ng.608 (2010).

108 Yang, J., Lee, S. H., Goddard, M. E. & Visscher, P. M. GCTA: a tool for genome-wide complex trait analysis. Am J Hum Genet 88, 76–82, doi:10.1016/j.ajhg.2010.11.011 (2011).

109 Lee, S. H., Wray, N. R., Goddard, M. E. & Visscher, P. M. Estimating missing heritability for disease from genome-wide association studies. Am J Hum Genet 88, 294–305, doi:10.1016/j.ajhg.2011.02.002 (2011).

110 Lee, S. H., Yang, J., Goddard, M. E., Visscher, P. M. & Wray, N. R. Estimation of pleiotropy between complex diseases using single-nucleotide polymorphism-derived genomic relationships and restricted maximum likelihood. Bioinformatics 28, 2540–2542, doi:10.1093/bioinformatics/bts474 (2012).

111 Bulik-Sullivan, B. K. et al. LD Score regression distinguishes confounding from polygenicity in genome-wide association studies. Nat Genet 47, 291–295, doi:10.1038/ng.3211 (2015).

112 Chang, C. C. et al. Second-generation PUNK: rising to the challenge of larger and richer datasets. Gigascience 4, 7, doi:10.1186/s13742-015-0047-8 (2015).

113 Plink v1.9 (2015).

114 Sun, L., Craiu, R. V., Paterson, A. D. & Bull, S. B. Stratified false discovery control for large-scale hypothesis testing with application to genome-wide association studies. Genet Epidemiol 30, 519–530, doi:10.1002/gepi.20164 (2006).

115 Storey, J. D. & Tibshirani, R. Statistical significance for genomewide studies. Proc Natl AcadSciUSA 100, 9440–9445, doi:10.1073/pnas.1530509100 (2003).

116 Consortium, G. T. Human genomics. The Genotype-Tissue Expression (GTEx) pilot analysis: multitissue gene regulation in humans. Science 348, 648–660, doi:10.1126/science. 1262110 (2015).

117 Pers, T. H. et al. Biological interpretation of genome-wide association studies using predicted gene functions. Nat Commun 6, 5890, doi:10.1038/ncomms6890 (2015).

118 Roadmap Epigenomics, C. et al. Integrative analysis of 111 reference human epigenomes. Nature 518, 317–330, doi:10.1038/nature14248 (2015).

119 Zambon, A. C. et al. GO-Elite: a flexible solution for pathway and ontology over-representation. Bioinformatics 28, 2209–2210, doi:10.1093/bioinformatics/bts366 (2012).

120 Kasprzyk, A. et al. EnsMart: a generic system for fast and flexible access to biological data. Genome Res 14,160–169, doi:10.1101/gr.1645104 (2004).

121 Kang, H. J. et al. Spatio-temporal transcriptome of the human brain. Nature 478, 483–489, doi:10.1038/nature10523 (2011).

